# Long distance avian migrants fail to bring 2.3.4.4b HPAI H5N1 into Australia for a second year in a row

**DOI:** 10.1101/2024.03.06.583767

**Authors:** Michelle Wille, Robyn Atkinson, Ian G. Barr, Charlotte Burgoyne, Alexander L. Bond, David Boyle, Maureen Christie, Meagan Dewar, Tegan Douglas, Teagan Fitzwater, Chris Hassell, Roz Jessop, Hiske Klaassen, Jennifer L. Lavers, Katherine K.-S. Leung, Jeremy Ringma, Duncan R. Sutherland, Marcel Klaassen

## Abstract

There is an ongoing and profound burden of lineage 2.3.4.4b high pathogenicity avian influenza (HPAI) H5N1 on wildlife and poultry, globally. Herein we report the continued absence of HPAI and antibodies against lineage 2.3.4.4b HPAI from October – December 2023, in migratory birds shortly after their arrival in Australia. Given the ever-changing phenotype of this virus, worldwide studies on the occurrence, or here absence of the virus, are of critical importance to understand the virus’ dispersal and incursion risk and development of response strategies.

## Main Text

The current high pathogenicity avian influenza (HPAI) H5N1 lineage 2.3.4.4b panzootic is having a profound impact on the poultry industry and wildlife (1, 2). HPAI H5N1 emerged in poultry in 1996 and has caused outbreaks in wild bird populations episodically since 2005 (3), with the epidemiology of the virus changing substantially with the emergence of new lineages. A novel lineage emerged in 2014 (2.3.4.4), which has diversified and caused substantial mortality, including mass mortality events, of wild birds in 2014, 2016, 2020-present, marine mammals since 2023, as well as ongoing outbreaks in poultry, globally (3). Understanding the changing phenotype and viral incursion risk following the emergence of novel lineages of HPAI is of crucial importance for the development of short-term and long-term mitigation strategies to protect wildlife, livestock and humans alike.

Wild birds, particularly waterfowl were initially implicated in the long distance spread of HPAI and have been predominantly implicated in the re-occurring incursions into Europe and Africa (4). However, recent viral incursions, such as that to the sub-Antarctic islands, and expansion, such as the movement of HPAI down the spine of South America (5) were driven by seabirds, suggesting that the long-distance dispersal of lineage 2.3.4.4b HPAI is no longer waterfowl dependant [*e*.*g*. (2, 6)].

Lineage 2.3.4.4b has now been detected on all continents except Oceania (7). HPAI incursion risk to Australia has previously been considered low due to the absence of waterfowl species that migrate beyond the Australio-Papuan region (8), and from influenza genomic surveillance (9). In turn, annually, millions of migratory seabirds and shorebirds migrate from Asia and North America to Australia (Figure 1). Some of these species have been shown to be part of the avian influenza reservoir community (10), have been infected by HPAI, and potentially survive and move HPAI viruses (11).

**Figure 1.**
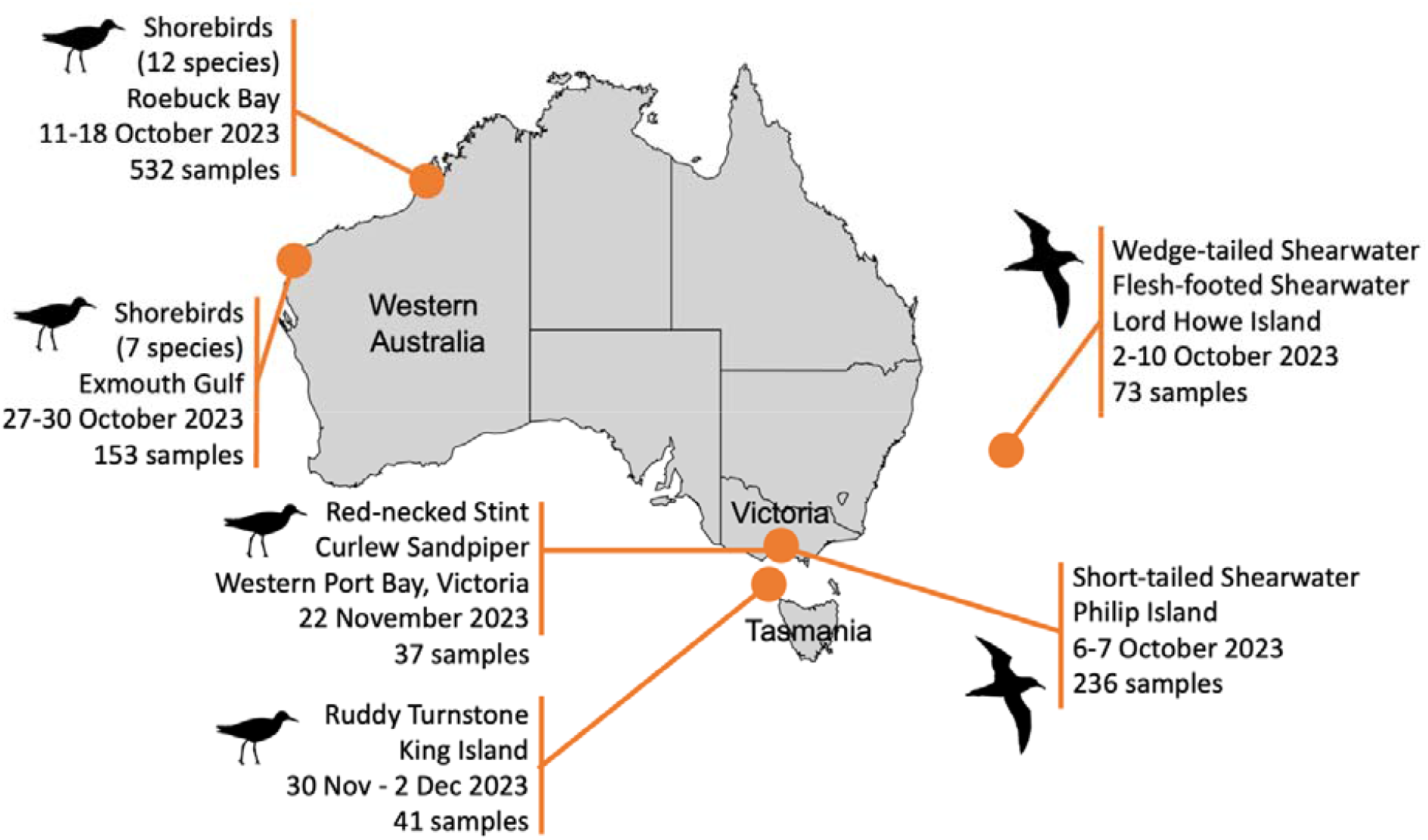
Sampling of incoming migratory birds in Australia for HPAI. All samples were negative for influenza A virus. Birds silhouettes from phylopic.com, distributed under a creative commons attribution.

To reveal whether a viral incursion may have occurred in Australia in 2023 with the arrival of wild migratory sea- and shorebirds, we investigated 1072 migratory birds of the order *Charadriiformes* and *Procellariiformes*, in October–December 2023. Specifically, we captured and sampled Short-tailed shearwaters (n=236) at a breeding colony on Phillip Island, Victoria, and Wedge-tailed (n=30) and Flesh-footed shearwaters (n=43) at a breeding colony on Lord Howe Island, New South Wales, upon their arrival from the northern Pacific. We also sampled twelve Asian-breeding migratory shorebird species at major non-breeding sites in Roebuck Bay (n=532) and Exmouth Gulf (n=153), in Western Australia, Western Port Bay in Victoria (n=37) and on King Island, Tasmania (n=41) (Table 1, Figure 1). Capture, banding and sampling was conducted under ABBBS authorities 2824 to Tegan Douglas, 2915 to MK, 8000 to Australasian Wader Studies Group and 8001 to the Victorian Wader Study Group, approval of animal ethics committees of Deakin University (B39-2019), Natural Resources and Environment Tasmania (AEC Project No. 6/2022-23 NRE), Philip Island Nature Parks (SPFL20082), Department of Primary Industries and Regional Development Western Australia (WAEC 23-08-52), Charles Stuary University (A22382).

**Table 1.**
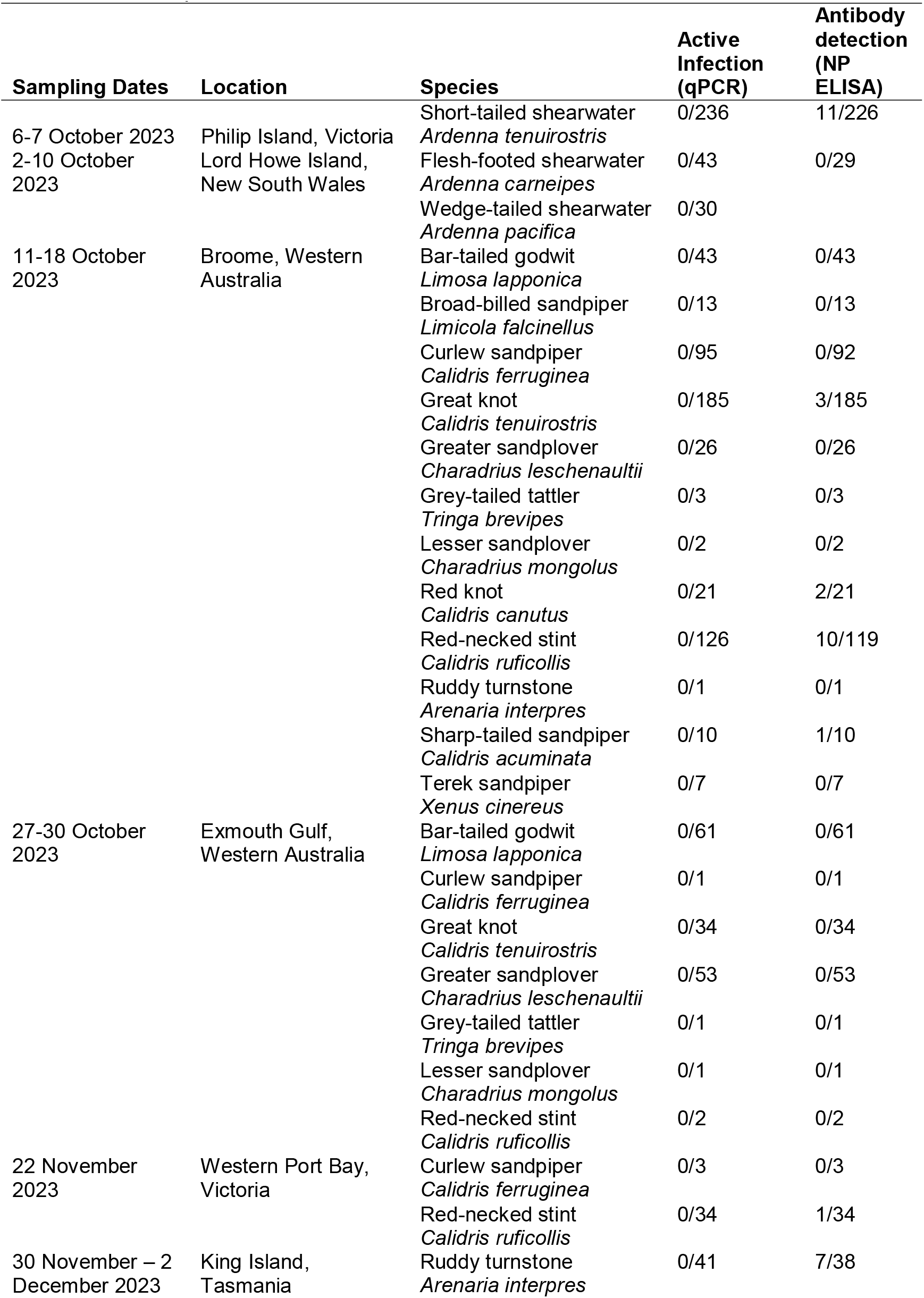
Charadriiform and Procellariiform species targeted for active HPAI surveillance following arrival to Australia after migration from Asia and North America. The number of active infections with influenza A virus and number of individuals with nuclear-protein antibodies against influenza A virus are depicted, compared with total number of individuals sampled.

All samples were negative for influenza A virus by qPCR, following (10). Thirty-six serum samples tested positive for anti-NP antibodies using a commercial ELISA (given an S/N cut off of 0.6)(12) (Table 1), which fell within the previously reported seroprevalence of the species that tested positive: Great knot, Red knot, Red-necked stint, Ruddy turnstone, Sharp-tailed sandpiper, and Short-tailed shearwater (10). Thirty-two sera samples positive by anti-NP ELISA which had sufficient sample volume remaining were subjected to a hemagglutination inhibition (HI) assay using lineage 2.3.4.4b virus A/Astrakhan/3212/2020(H5N8) and A/American wigeon/South Carolina/22-000345-001/2021(H5N1), as well as a Australian lineage LPAI H5N3 Australian lineage virus A/duck/Victoria/0305-2/2012(H5N3) following (11). Both 2.3.4.4b antigens were 6:2 recombinant virus on an A/Puerto Rico/8/1934(H1N1)(PR8) backbone with the multi-basic cleavage site removed. None of the 32 serum samples tested had evidence for anti-HA antibodies against either 2.3.4.4b antigen, and one sample (17696) from a Short-tailed Shearwater had anti-HA antibodies against the LPAI H5 Australian lineage virus.

In cases where wild bird mortality events occurred in Australia during this period, all tested birds were negative for HPAI (pers. comm. Wildlife Health Australia).

For Australia, as for other regions in the world, HPAI incursion risk hinges on a combination of factors, including wild bird migration, virus pathogenicity in migratory birds (notably whether birds can migrate while infected), and virus circulation in neighbouring regions. That, like in 2022 (13), there was no incursion of HPAI in Australia in 2023, despite the arrival of millions of migratory birds, could possibly be attributed to two factors. Firstly, while the virus’ host reservoir has substantially widened and intensified beyond waterfowl (2, 6, 14), waterfowl are markedly absent from the list of migrants to Australia.

HPAI infected waterfowl appear to migrate without negative effects, but it is unclear whether migratory culling may prevent the arrival of HPAI infected shorebirds and seabirds to Australia. Secondly, migrants to Australia predominantly use the East Asian Australasian Flyway (EAAF). Aside from few countries, such as Japan, there are few HPAI outbreaks reported in wild birds along the EAAF (2), suggesting a potentially low virus exposure potential for long distance migratory birds using the EAAF. Further, in Asia, ancestral HPAI H5 lineages, such as 2.3.2.1c in Cambodia or 2.3.2.1a in Bangladesh (3), are endemic and continue to circulate. Exposure to these more ancestral lineages may potentially result in cross-protection against 2.3.4.4.b in both poultry and wild birds, serving as a potential explanation why outbreaks of lineage 2.3.4.4b HPAI are remarkably limited in East Asia (2). None of the migrants tested in this study and in our 2022 study (13) had antibodies against lineage 2.3.4.4b HPAI, and no incursion of lineage 2.3.4.4b HPAI to Australian continent has occurred. These two hypotheses warrant further investigation.

Australia will again enter a high-risk period when the major bird migrations into the country take place between August - November 2024. Continued surveillance is critical for early detection and rapid response, and as such we call for enhanced surveillance of Australian wild birds to match heightened incursion risk in the second half of 2024.

## Acknowledgements

We wish to acknowledge staff, members and volunteers associated with the Australasian Wader Studies Group, Birdlife Australia, Philip Island Nature Parks, Victorian Ornithological Research Group, and Victorian Wader Study Group, who contributed to bird capture and sample collection, and K. Milne for sample shipping support. We are grateful to T. Grillo, S. Ban, S. Vitali and A. Breed for ongoing support.

This work was funded by the Department of Agriculture, Fisheries and Forestry One Health Investigation Fund (administered by Wildlife Health Australia), and Australian Research Council (ARC) Discovery Project Grant DP190101861 (to MK). The WHO Collaborating Centre for Reference and Research on Influenza is funded by the Australian Department of Health. CH and KKSL were funded by Beijing Normal and Princeton Universities, ALB is funded by the Natural History Museum. Exmouth samples were collected with the support of Woodside Energy.

We acknowledge the Yawuru People via the offices of Nyamba Buru Yawuru Limited for permission to catch birds on the shores of Roebuck Bay, traditional lands of the Yawuru people.

## Notes

### Competing Interest Statement

The authors have declared no competing interest.

